# Dynamic brain interactions during picture naming

**DOI:** 10.1101/478495

**Authors:** Aram Giahi Saravani, Kiefer J. Forseth, Nitin Tandon, Xaq Pitkow

**Author notes:** Corresponding authors *Email addresses:* (Aram Giahi Saravani), (Nitin Tandon), (Xaq Pitkow).

## Abstract

Brain computations involve multiple processes by which sensory information is encoded and transformed to drive behavior. These computations are thought to be mediated by dynamic interactions between populations of neurons. Here we demonstrate that human brains exhibit a reliable sequence of neural interactions during speech production. We use an autoregressive hidden Markov model to identify dynamical network states exhibited by electrocorticographic signals recorded from human neurosurgical patients. Our method resolves dynamic latent network states on a trial-by-trial basis. We characterize individual network states according to the patterns of directional information flow between cortical regions of interest. These network states occur consistently and in a specific, interpretable sequence across trials and subjects: a fixed-length visual processing state is followed by a variable-length language state, and then by a terminal articulation state. This empirical evidence validates classical psycholinguistic theories that have posited such intermediate states during speaking. It further reveals these state dynamics are not localized to one brain area or one sequence of areas, but are instead a network phenomenon.

## 1. Introduction

Neural computation requires the orchestration of distributed cortical processes. In many complex cognitive tasks, it is unlikely that interactions between large populations of neurons merely generate a simple feedforward sequence of activated brain regions. Instead, these processes likely involve distributed interactions that change over time and depend on context. To understand information flow and computation in the brain, it is critical to account for these dynamic interactions between functionally distinct regions.

For simplicity, most analyses of coarse-grained brain activity like that measured by electrocorticography (ECoG) are based on assumptions of linearity. These include methods based on second-order correlations (Aertsen et al., 1987; Friston et al., 1993), structural equation modeling (Horwitz, 1994; Penny et al., 2004), Granger causality (Granger, 1969; Baccalá and Sameshima, 2001; Josic et al., 2009), Gaussian graphical models (Zheng and Rajapakse, 2006; Rajapakse and Zhou, 2007), and linear dynamical systems. Such linear models cannot capture crucial behaviors like flexible time-dependent or context-dependent interactions. One way to improve the expressiveness of these models while preserving some of their tractability is to use a switching linear dynamical system, where the switch determines which linear model currently best describes the neural dynamics.

Here we present the first application of such a switching linear dynamical system to direct recordings from human brains during language production. We applied an autoregressive hidden Markov model (ARHMM), a type of hierarchical Bayesian network, that accounts for the observed continuous time series as a consequence of switching between a discrete set of network states that govern the electrical activity. Each discrete state corresponds to distinct stochastic linear dynamics for the observed recordings. These switching dynamics between the different network states approximates the nonlinear dynamics of the full system. Our statistical method learns these state dynamics and the latent state transition matrix, as well as the trial-specific state sequences (Section 2.6).

Recent studies of human ECoG data have used ARHMMs to classify dynamical modes of epileptic activity (Wulsin et al., 2014; Baldassano et al., 2016), a neural state characterized by relatively simple dynamics in comparison to healthy brain function. In the present work, we use ARHMMs to classify dynamical states and demonstrate their ability to reveal information flow between brain areas during normal cognition. In particular, we demonstrate that ARHMMs can identify and characterize transient network states by analyzing ECoG data recorded during picture naming, a uniquely human ability which involves visual processing, lexical semantic activation and selection, phonological encoding, and articulation.

Picture naming is an oft studied language task that has been the backbone of many psycholinguistic theories of speech production. Chronometric studies of picture naming (Levelt, 1989), as well as the nature of common speech errors (Fromkin, 1973) and speech disruption patterns in aphasia (Dell et al., 1997), have led to theories that linguistic components are organized hierarchically and assembled sequentially. Yet there are no data that can directly be used to validate these ideas, and the dynamics of the cortical networks supporting even simple, single-word articulations, remain unknown. In the absence of such data, it is difficult to resolve competing models such as between discrete (Fromkin, 1973; Garrett, 1980; Indefrey and Levelt, 2004) and interactive (Dell et al., 1997; Rapp and Goldrick, 2000) models of language production.

Prior studies have leveraged the high spatiotemporal resolution of intracranial electroencephalography to study specific brain regions during language production (Sahin et al., 2009; Edwards et al., 2010; Conner et al., 2014; Kadipasaoglu et al., 2016; Forseth et al., 2018), and have applied adaptive multivariate autoregressive (AMVAR) analysis (Ding et al., 2000; Korzeniewska et al., 2008; Whaley et al., 2016), to reveal the fast, transient dynamics of human cortical networks. This class of methods assumes a consistent progression of state sequences across trials, an assumption that is frequently violated during complex behaviors like language that invoke multiple cognitive processes. Consequently, the inferred activity patterns from AMVAR may be grouped into false network states. The ARHMM analysis developed here is a principled probabilistic framework to resolve trial-by-trial network state dynamics. As an added benefit, it also provides model uncertainties. We demonstrate that this method delivers improved estimates of network dynamics in human language function compared to conventional AMVAR clustering analyses.

We show that unsupervised Bayesian methods can infer reliable time series of latent network states and information flow from ECoG signals during a task requiring integration of visual, semantic, phonological, and sensorimotor processing. These states have dynamics that are consistent across subjects and reflect the timing of subjects’ actions. From the trial-resolved sequence network states we learn additional characteristics of the network states not available through fixed-timing models like AMVAR.

## 2. Materials and Methods

### 2.1. Human Subjects

We enrolled 3 patients (1 male, 2 female; mean age 28 ± 9 years; mean IQ 86 ± 3) undergoing evaluation of intractable epilepsy with subdural grid electrodes (left, n = 2; right, n = 1) in this study after obtaining informed consent. Study design was approved by the committee for the protection of human subjects at the University of Texas Health Science Center. Hemispheric language dominance was evaluated by intra-carotid sodium amytal injection (Wada and Rasmussen, 2007), fMRI laterality index (Ellmore et al., 2010; Conner et al., 2011), or cortical stimulation mapping (Tandon, 2008; Forseth et al., 2018). All patients were found to have left-hemisphere language dominance.

### 2.2. Experimental Design

Subjects engaged in a visual naming task (Figure 1D, left). We instructed subjects to articulate the name for common objects depicted by line drawings (Snodgrass and Vanderwart, 1980; Kaplan E, Goodglass H, 1983). Subjects were instructed to report “scrambled” for control images in which we randomly rotated pixel blocks demarcated by an overlaid grid (Figure 1D, right). Each visual stimulus was displayed on a 15-in LCD screen positioned at eye-level for 2 seconds with an inter-stimulus interval of 3 seconds. A minimum of 240 images and 60 scrambled stimuli were presented to each patient using presentation software (Python v2.7).

**Figure 1:**
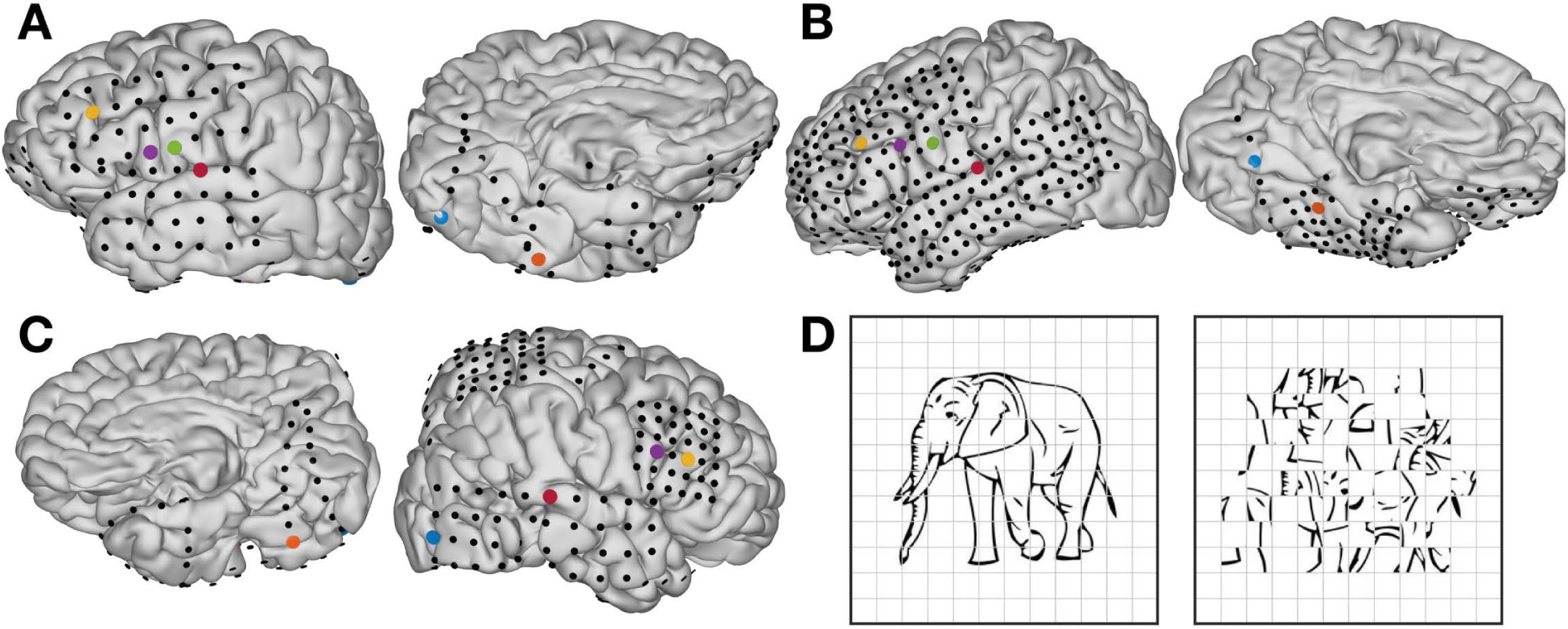
A-C, Individual pial surface and electrode reconstructions. One representative electrode was selected from each of the following regions: early visual cortex (blue), middle fusiform gyrus (orange), pars triangularis (yellow), pars opercularis (purple), ventral sensorimotor cortex (green), and auditory cortex (red). D, Picture naming stimuli: coherent (left) and scrambled (right).

### 2.3. MR Acquisition

Pre-operative anatomical MRI scans were obtained using a 3T whole-body MR scanner (Philips Medical Systems) fitted with a 16-channel SENSE head coil. Images were collected using a magnetization-prepared 180^*◦*^ radiofrequency pulse and rapid gradient-echo sequence with 1 mm sagittal slices and an in-plane resolution of 0.938 × 0.938 mm (Conner et al., 2011). Pial surface reconstructions were computed with FreeSurfer (v5.1) (Dale et al., 1999) and imported to AFNI (Cox, 1996). Post-operative CT scans were registered to the pre-operative MRI scans to localize electrodes relative to cortex. Subdural electrode coordinates were determined by a recursive grid partitioning technique and then validated using intra-operative photographs (Pieters et al., 2013).

### 2.4. ECoG Acquisition

Grid electrodes (n = 615) — subdural platinum-iridium electrodes embedded in a silastic sheet (PMT Corporation; top-hat design; 3 mm diameter cortical contact) — were surgically implanted via a craniotomy (Tandon, 2008; Conner et al., 2011; Pieters et al., 2013). ECoG recordings were performed at least two days after the craniotomy to allow for recovery from the anesthesia and narcotic medications. These data were collected at either a 1000 or 2000 Hz sampling rate and a 0.1–300 Hz or 0.1–700 Hz bandwidth, respectively, using NeuroPort NSP (Blackrock Microsystems). Continuous audio recordings of each patient were made with an omnidirectional microphone (30–20,000 Hz response, 73 dB SNR, Audio Technica U841A) placed next to the presentation laptop. These were analyzed offline to transcribe patient responses and to determine the time of articulation onset and offset (Forseth et al., 2018).

### 2.5. Data Processing

From previous work, we identified 6 anatomical regions of interest that broadly span the cortical network engaged during picture naming (Conner et al., 2014; Forseth et al., 2018): early visual cortex, mid-fusiform gyrus, pars triangularis, pars opercularis, ventral sensori-motor cortex, and superior temporal gyrus. In each individual, we selected an electrode with significant activity from each region (Figure 1A,B). These electrodes were uncontaminated by epileptic activity, artifacts, or electrical noise. Furthermore, we analyzed only trials in which cortical activity did not show evidence of epileptiform artifact (Conner et al., 2014; Kadipasaoglu et al., 2014; Forseth et al., 2018).

We selected trials with reaction times > 600 ms and < 1800 ms. Data was re-referenced to a common average of electrodes without epileptiform activity. Instantaneous gamma power (60–120 Hz)was extracted at each channel using a bandpass Hilbert transform. This absolute measure was then normalized to a pre-stimulus baseline (–700 ms to –200 ms relative to picture presentation).

Data were downsampled to 200 Hz. To initialize the ARHMM, we estimated the dynamics in 100 ms time windows with 50 ms overlap, following the AMVAR estimation method of (Ding et al., 2000). The AMVAR estimates were clustered (*k*-means clustering with Euclidean distance norm) into a set of discrete states to initialize the ARHMM inference.

### 2.6. Autoregressive Hidden Markov Model

Autoregressive (AR) processes are random processes with temporal structure, where the current state ***x**_t_* of a system is a linear combination of previous states and a stochastic innovation ***ν**_t_* ∼ 𝒩(0*, I*) (zero mean isotropic white noise). The dynamics of such a system is thus both linear and stochastic. This stochastic linear dynamical system can be described by a tensor of autoregressive coefficients, ***A*** = {*A_τ_*} (*i.e.* a matrix for each time lag *τ*), and a covariance matrix *Q* for the stochastic aspect of the system:

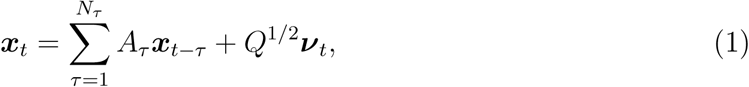

where *N_τ_* is the order (number of relevant time lags) of the autoregressive process. Since this model is linear, it is poorly suited to describing brain activity, which motivates us to use a richer model with changing dynamics.

A Hidden Markov Model (HMM) is a latent state model which describes observations as a consequence of unobserved discrete states *z*, where each state emits observable variables with specific probabilities. The probability of occurrence of a state depends on the previous state, the defining characteristic of a Markov model. The set of transition probabilities constitute the state transition matrix,

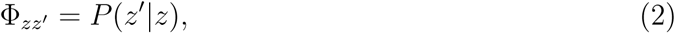

where *z′* and *z* are the current and previous state, respectively.

The ARHMM combines autoregressive stochastic linear dynamics with the Hidden Markov Model (Ghahramani and Hinton, 2000; Fox et al., 2009): Each latent state *z* indicates a different linear dynamical system with state-specific dynamics and process noise covariance, ***A**_z_* and *Q_z_* (Figure 2A,B). The switching of the linear dynamical systems makes the ARHMM effectively nonlinear.

**Figure 2:**
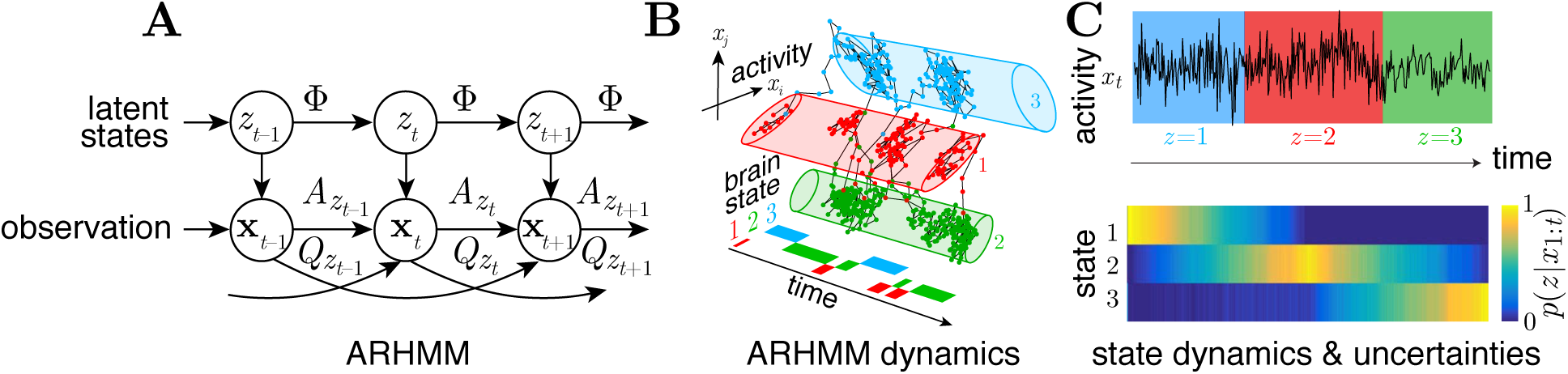
**A**. Graphical representation of an ARHMM with autoregressive order 2: latent states *z* and observations *x* evolve according to state transition matrix Φ, autoregressive coefficients ***A***, and process covariance *Q*. **B**. Illustration of the ARHMM latent state space model. **C**. Simulated time series of data points emitted by 3 latent states (top) with inferred state probabilities (bottom)

Applied to the multivariate ECoG data, each state describes the measured neural activity by a different linear dynamical system, with a set of AR coefficients, *A_zτij_*, that describe the (Granger) causal dynamical relationship between nodes *i* and *j* for state *z* at time lag *τ*. For a given state *z* at time *t*, the MVAR coefficients constitute an MVAR tensor, ***A**_z_t__*, describing the evolution of the multivariate ECoG signal ***x*** at time *t*,

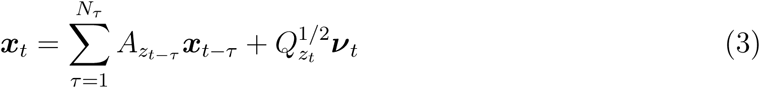

where ***ν*** 𝒩(0*, I*).

Neither the set of dynamical parameters, {***A***, *Q*}, nor the occurrence and frequency of individual states (represented by the state transition matrix Φ), are observed. The ARHMM infers the latent parameters: the time series of network states *z_t_*, their transition probabilities Φ_*zz*′_, and the dynamical parameters for each state, ***A**_z_* and *Q_z_*.

This is in contrast to the commonly used method in ECoG signal analysis to estimate the autoregression coefficient matrix ***A*** and process noise covariance, *Q*, based on the assumption that the network states occur at same times across all trials (Morf et al., 1978; Ding et al., 2000). This assumption will inevitably be inaccurate when there are significant variations in the emergence and durations of state sequences. This leads to artifacts in the estimation of the state parameters and the creation of pseudo-states that combine data points from different states into a new mixed state estimate. Also, our method does not require any manual alignment of trials by the epoch of interest, such as stimulus onset or articulation onset (Whaley et al., 2016), but instead provides a means of predicting these events from brain activity.

Probabilistic inference alleviates this problem by assigning posterior probabilities (conventionally called ‘responsibilities’), *P* (*z_t_│**x***_1*..T*_), to each state *z_t_* given the entire observed data sequence ***x***_1*..T*_ (Figure 2C,D). Since neither responsibilities nor dynamical parameters are known *a priori*, estimates for state parameters and responsibilities are calculated iteratively by an expectation-maximization algorithm (EM) (Dempster et al., 1977) known as the Baum-Welch algorithm (Baum et al., 1970). We incorporate a prior over Φ to favor infrequent transitions, by adding pseudocounts of self-transitions to the observed state transitions. This is realized by adding a scaled identity matrix, *I*, to Φ, and then renormalizing:

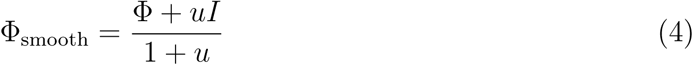

where *u* sets the lower bound on the time constant of self-transitions, and thereby determine minimal average state durations.

Initial conditions for ***A*** and *Q* are informed by the lagged correlation MVAR clustering method from (Morf et al., 1978; Ding et al., 2000). For the expectation part of our algorithm, responsibilities within each trial are evaluated based on the previous iteration’s parameters from the maximization loop. The maximization in turn uses these responsibilities to attribute the data points to different network states when estimating new parameters. The number of states and time lags in the ARHMM is selected according to the Bayesian Information Criterion (BIC) (Schwarz, 1978).

The ARHMM classifies dynamical states by the network connectivity associated with the inferred MVAR coefficient matrix. The MVAR coefficients and the related partial directed coherence in frequency domain are measures of causality for interactions between brain areas. Partial directed coherence (PDC) was defined by Baccalá and Sameshima (2001) to describe information flow (in the sense of Granger causality) between multivariate time series in the frequency domain. This measure is directly related to the MVAR coefficients, and for each state, *z*, we have:

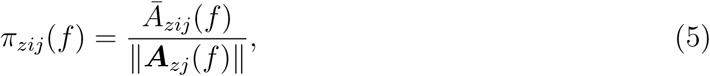

where

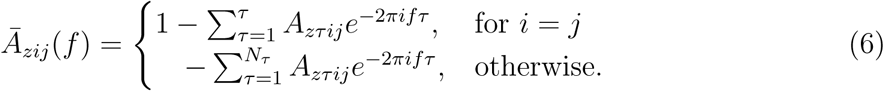

represents the transfer function at frequency *f*, 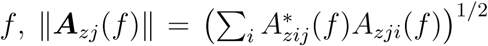 is the norm of the *j*th column of *A_z_*(*f*), and ^*^ denotes the complex conjugate. PDC is normalized to show the ratio between the outflow from channel *j* to channel *i* to the overall outflow from channel *j*, taking values from the interval [0, 1].

### 2.7. Visualizing network states

For each latent state *z*, we have an associated set of MVAR coefficients ***A**_z_*. We display these as directed graphs, with interactions quantified by PDC integrated over all frequencies. In these graphs, each node represents a single electrode and arrows represent the causal relationship between nodes.

To visualize the inferred time series of network states with their associated probability we display the statistically most likely sequence of states (Viterbi trace) weighted by the associated uncertainty (responsibility).

### 2.8. Robustness of ARHMM inference

To see how robust the ARHMM is to deviations from our model assumptions, we simulated data with mean-dependent process noise variance and observation noise. Note that uncontrolled or unmodeled factors can have similar effects as observation noise, creating unexplained variability. Figure 3A shows the simulated multivariate signal and its mean-dependent variance. Initialized from AMVAR-based *k*-means clustering of network states, the ARHMM then infers the trial-by-trial state sequence (Figure 3B). Depending on the amount of observation noise (signal-to-noise ratios of SNR=0.3, 3, and 30), the inference has different degrees of confidence, visualized by the brightness. (Figure 3C) shows network interactions for each state, again for different signal-to-noise ratios. The inferred states and network properties in each state generally agree with the ground truth despite the model mismatch from observation noise and mean-dependent variance. The beginning and the end of the trials have similar properties, and for lower signal-to-noise ratios (more observation noise) the ARHMM naturally identifies them with the same latent state. Interestingly, in these conditions the ARHMM also misclassifies states in the middle of the trials, as it auto-matically finds hallmarks of the first state in the middle of the trial. For smaller observation noise, these states are correctly classified. Even when some times are misclassified in low SNR, the network structures are recovered accurately (Figure 3C).

**Figure 3:**
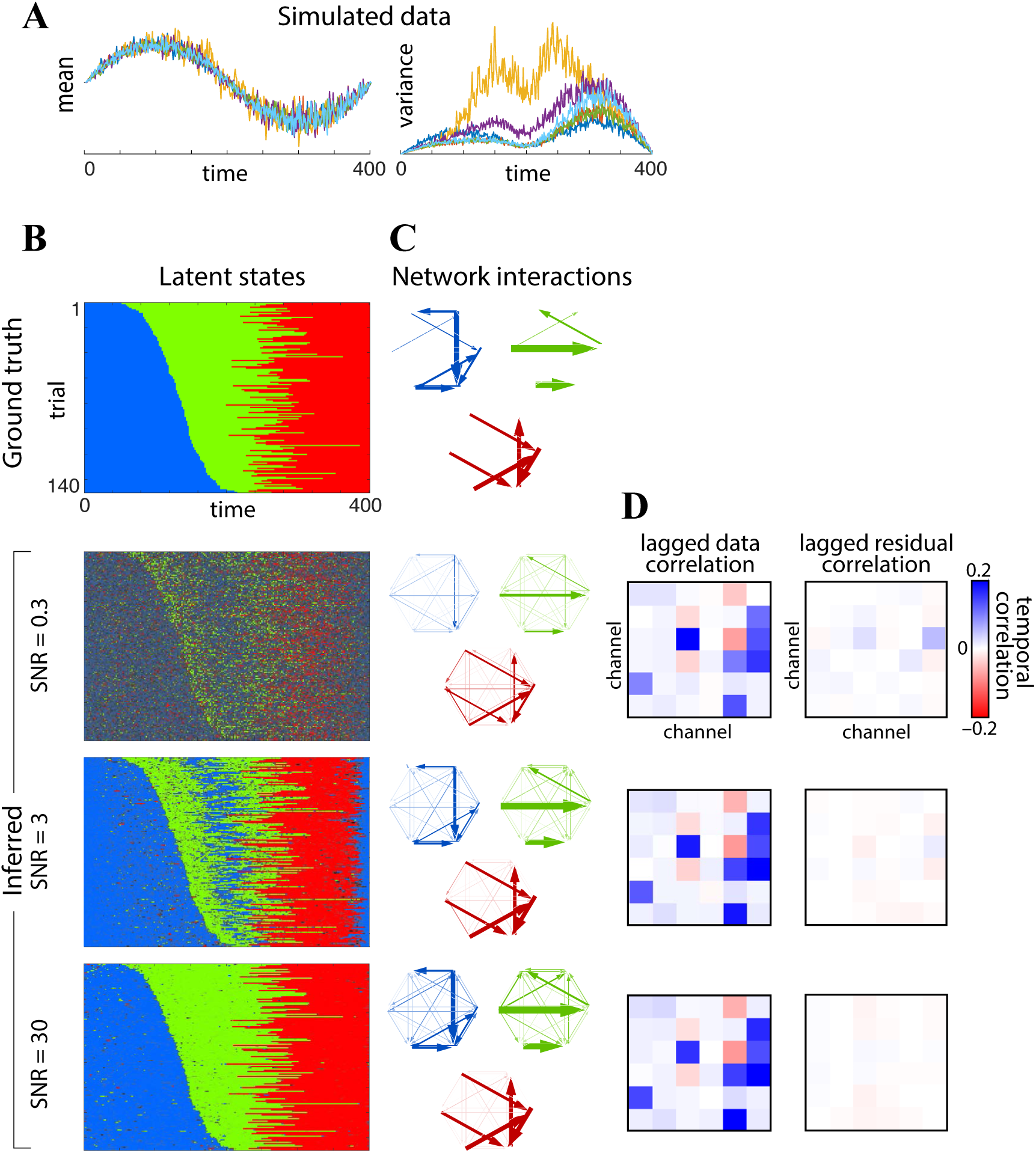
Comparison of state clustering with different data pre-processing methods for simulated data. (**A**) Underlying data with varying state durations, and mean-dependent variance with associated state sequence and network states on the right. (**B**) Rasters of discrete latent states over time and trials, for ground truth (top panel) and estimates (bottom three panels) inferred from data with three different signal-to-noise ratios (SNR=0.3, 3, and 30). Estimates are obtained from mean-subtracted time series. Color represents the discrete states, and confidence (responsibility) is encoded by brightness and increases with SNR. (**C**) Graphs of network interactions for each latent state, measured by Partial Directed Coherence (PDC). (**D**) Spatial correlation structure at a time lag of 1 for measured signal (left) and residuals between ARHMM fit and data (right).

To estimate how well our model explains the temporal structure of the data we follow (Ding et al., 2000) and perform a residual whiteness test for our model fit, computing the auto- and cross-correlation of the multivariate residuals between the ARHMM fit and the data for each time lag up to AR model order (except zero), as shown in Fig. 3D. If the model captures the temporal structure of the data, the auto- and cross-correlation coefficients of the residuals should approach zero (uncorrelated white noise). The residuals are indeed small, indicating that the model is reasonable even for small SNR.

### 2.9. Statistical Analysis

Correlations in Figure 6 are calculated using the Pearson correlation. Significance was evaluated using a two-sided test for deviations from zero correlation. This was evaluated using the Fisher transform, which renders correlations approximates normal and thereby gives *p*-values as 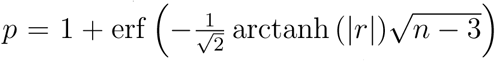 where *n* is the number of samples and *r* is the measured correlation coefficient (Fisher, 2006).

## 3. Results

We observed a sequence of peak trial-averaged GBA (Fig. 4A) beginning in early visual cortex, moving anteriorly to middle fusiform gyrus and pars triangularis, and then culminating in pars opercularis, subcentral gyrus, and superior temporal gyrus. Consistent with prior work (Conner et al., 2014; Forseth et al., 2018), Fig. 4B indicates that the GBA response varies substantially across trials. This motivates an analysis method like the ARHMM that can account for the such variability.

**Figure 4:**
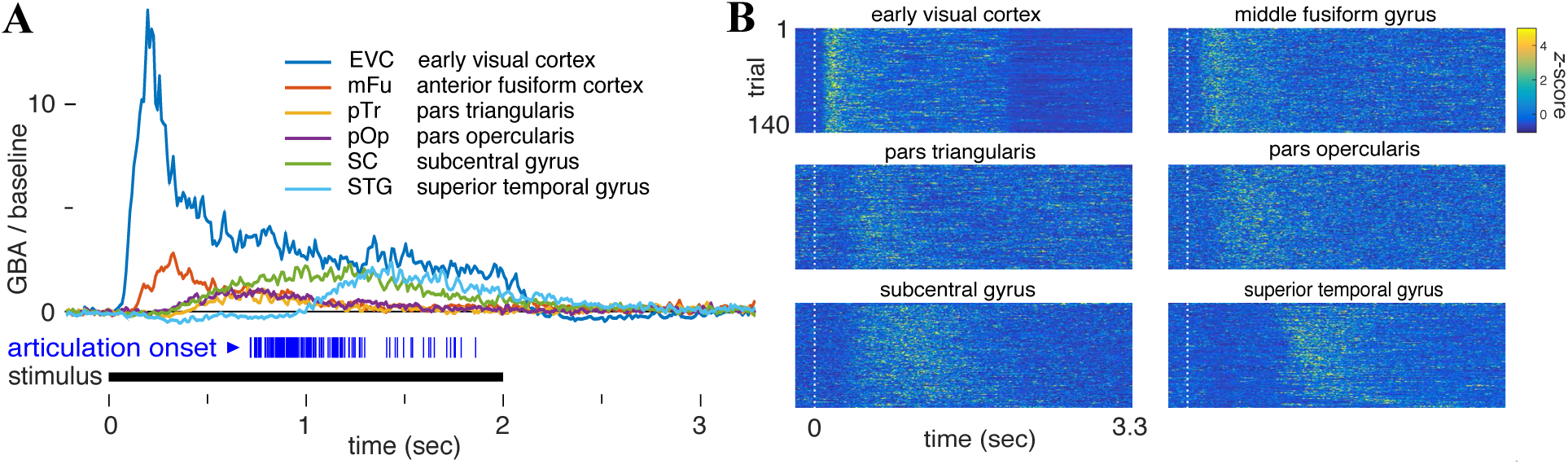
Gamma-band activity (GBA) at salient brain regions following picture presentation. **A**: Trial-averaged GBA, relative to baseline. Each articulation onset is indicated by a vertical blue line below (mean 1.2 sec), and the visual stimulus is presented during the black interval. **B**: Density plot of trial-wise GBA, *z*-scored.

Using ARHMM inference, we analyzed the network dynamics for each subject independently (Fig. 5). These analyses converged on models for all subjects featuring dynamics that depended on three time lags and were best explained by four network states. Pre-stimulus and post-articulation activity patterns were predominantly assigned to one state, which we therefore named ‘resting’. Immediately following picture presentation, a second state dominated for ~250 ms, which we named ‘visual processing’. A prominent feature of this network state was information flow from early visual cortex and middle fusiform gyrus. The next state is characterized by interactions distributed throughout the network, but most strongly driven by frontal regions. We named this state ‘language processing’. Comparing our inference results for both hemispheres, we find that the ‘language processing’ state was only pronounced in the recordings of language-dominant cortex (Fig. 5), consistent with a left-lateralized language production network. During articulation, we observed a fourth state we named ‘articulation’ which featured greater information flow from subcentral gyrus and superior temporal gyrus back to the language processing network. The ARHMM model reveals that neural interactions transition back to the ‘resting’ state following each completed articulation.

**Figure 5:**
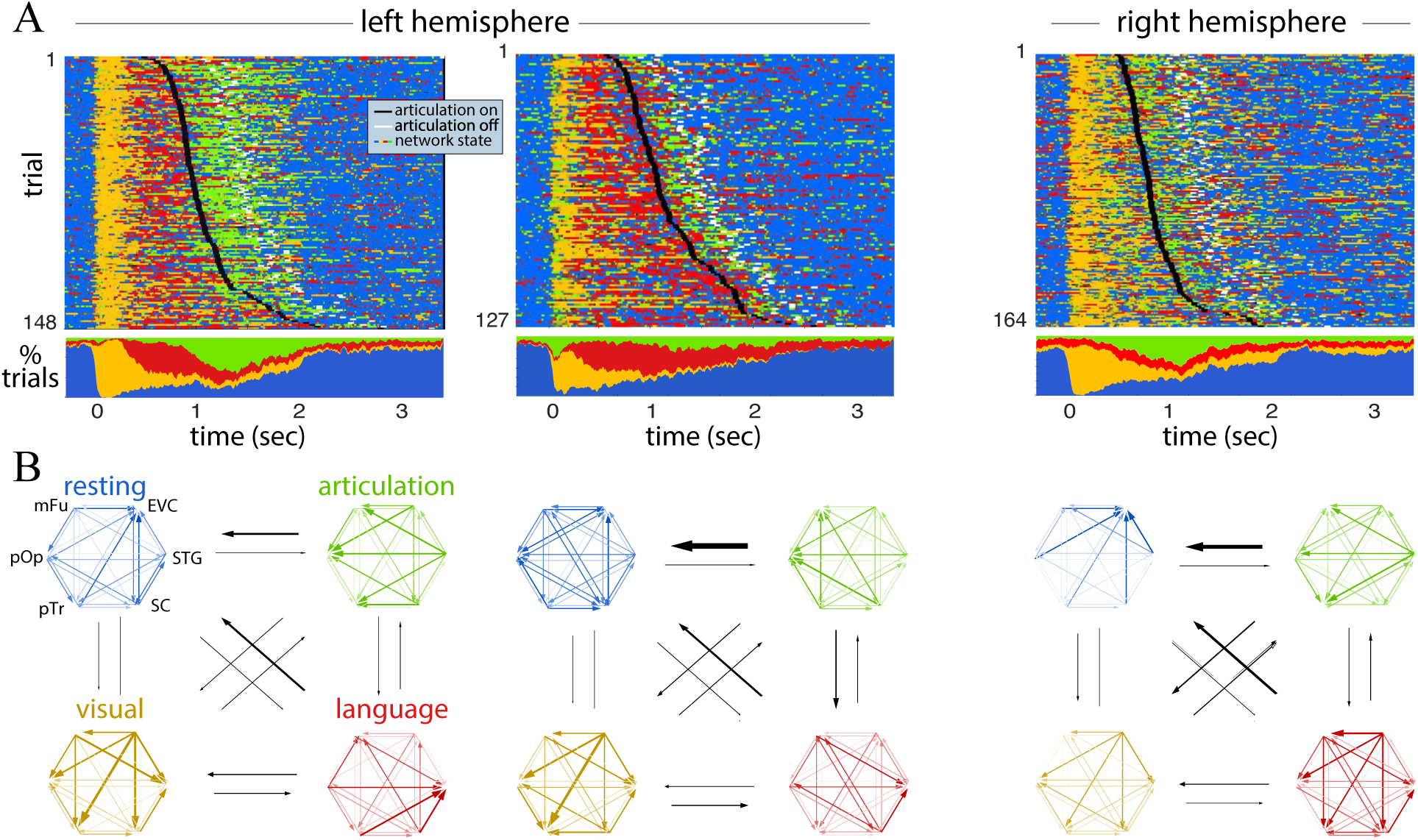
State sequences in brain activity. **A**: Time dependence of most probable states from left and right hemispheric brain activity for three subjects. The top shows states as a trial-by-trial raster, and the bottom shows the fraction of trials on which each state is most probable. **B**: Interactions between brain regions during the corresponding named network states, plotted as in Fig. 3C. Black arrows indicate state transition probabilities according to the inferred state transition matrix.

**Figure 6:**
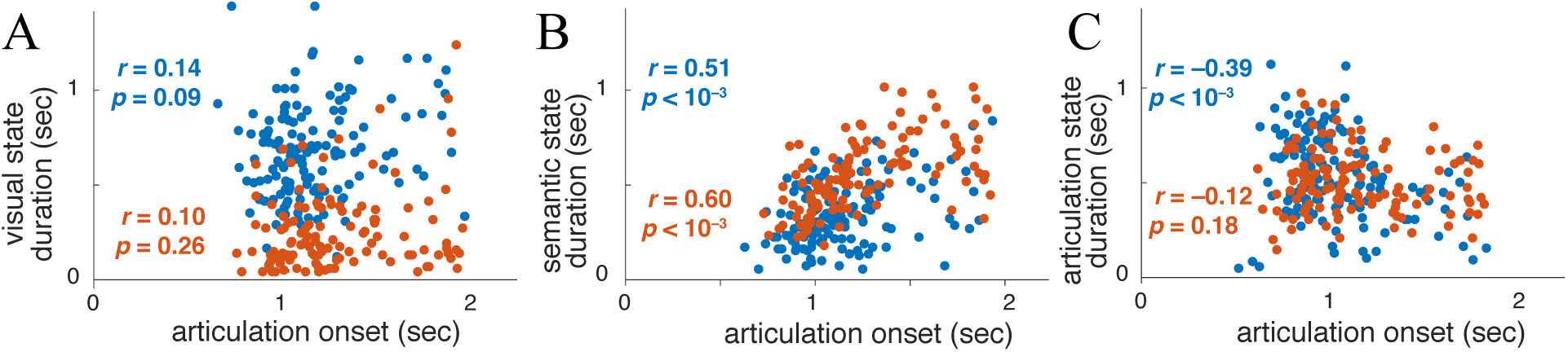
Duration of the active states, (**A**) ‘visual processing’, (**B**) ‘language processing’, and (**C**) ‘articulation’, for two patients’ left-hemispheres (blue and orange), compared to reaction times. The Pearson correlation coefficient *r* and *p*-value for the null hypothesis of uncorrelated values are shown.

An indication that the network states found by the ARHMM reflect task-relevant neural computations is that they predict the onset of articulation. The duration of the ‘visual processing’ state was uninformative about this onset time (Fig. 6A), but the duration of the ‘language processing’ state was significantly related to reaction time (Fig. 6B). This is consistent with variable difficulty in identifying word names for the heterogeneous stimuli. For one of the two patients with left hemisphere recordings, the duration of the articulation state was *negatively* related to articulatory onset (Fig. 6C).

Across these three patients and across 120–160 trials, we find that the inferred network states follow a reliable state sequence (Fig. 5A). Moreover, each state’s interactions between brain regions were similar between patients, as measured by partial directed coherence (PDC) (Fig. 5B). Our measurements do not cover the complete visual and language processing networks, and these areas are unlikely to have exclusively direct connections. Nonetheless, the analysis still indicates that interactions between early visual cortex and the language areas have directionality, even if mediated by some unmeasured areas.

The control condition of scrambled images still provides a visual stimulus and requires articulation, but does not bind a specific concept (aside from the general category of “scrambled”). We measured differences in network dynamics during coherent and scrambled image naming by inferring the network structure and states for each condition separately. To improve the ARHMM inference and to make it more robust with respect to subject-to-subject variability, we combined trials from both left-hemisphere data sets, effectively assuming that electrodes in both recordings were anatomically homologous.

Neural activity in both stimulus conditions were described by comparable networks, but exhibited specific differences in connectivity between the network nodes (Fig. 7 left). The differences were most pronounced in the active states: ‘visual processing’, ‘language processing’, and ‘articulation’. This suggests that the observed differences are due to condition-specific network activity.

**Figure 7:**
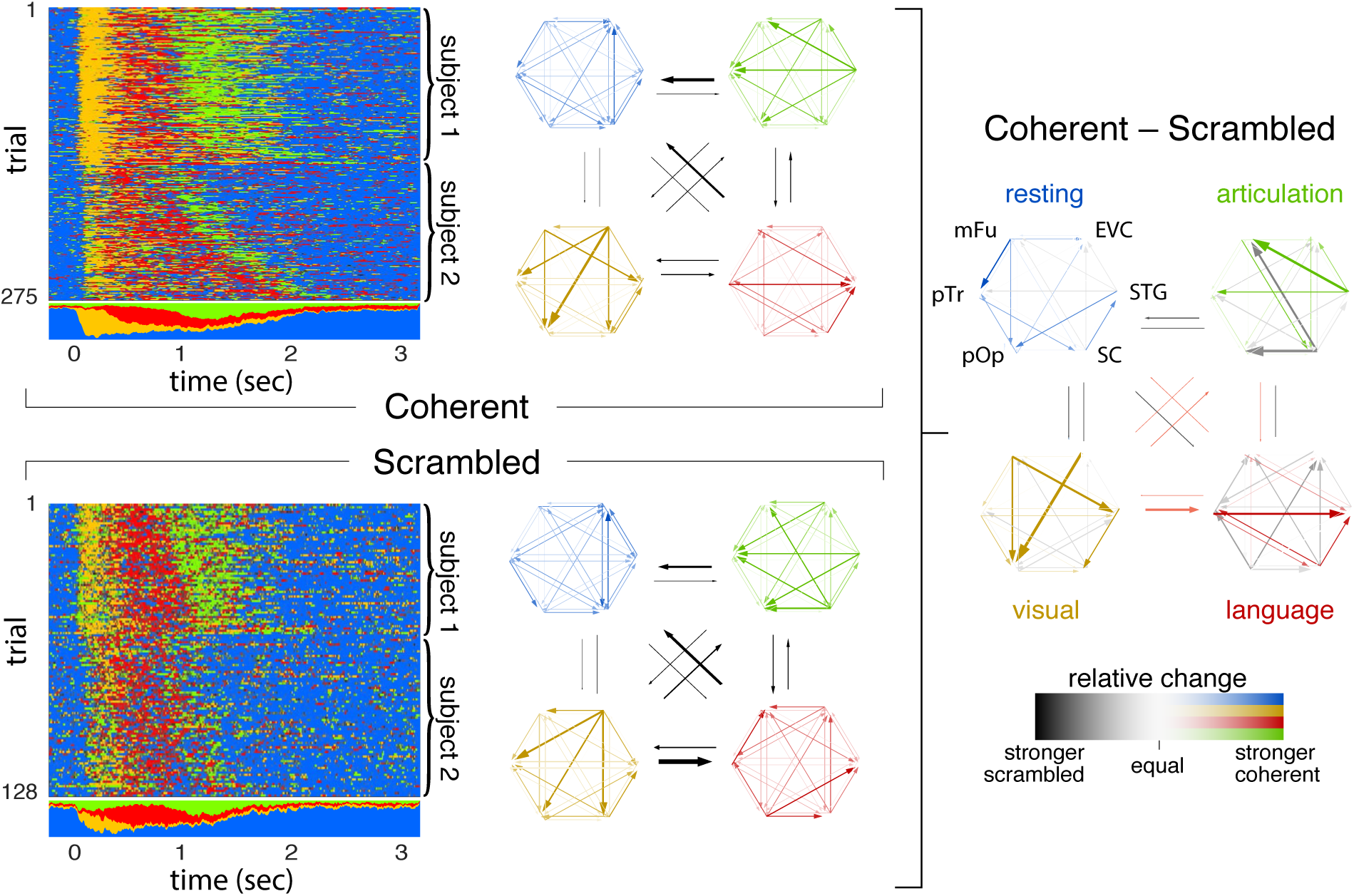
Multi-subject integration (anatomical grouping) of the two left-hemispheric recordings for coherent (top left) and scrambled (bottom left) stimulus condition. Differences in information flow between both stimulus conditions are shown on the right, where colored and gray arrows denote excess activity in the coherent and scrambled condition, respectively.

The strongest network connections in each state were generally suppressed when viewing scrambled images. During the ‘visual processing’ state for coherent images, early visual cortex and middle fusiform gyrus had stronger influences on frontal regions than for scrambled images. In the ‘language processing’ state, the superior temporal gyrus received more input from subcentral gyrus and Broca’s area (pars triangularis and pars opercularis) when naming coherent images. The ‘articulation’ state for coherent images shows increased information flow emanating from superior temporal gyrus, while the same state for scrambled images shows increased information flow from subcentral gyrus. This dissociation between auditory and sensorimotor cortex responses to coherent and scrambled images could be driven by learning effects from consistent repetition of the stereotyped response “scrambled.”

As described above, we find that Broca’s area — pars triangularis and pars opercularis — strongly interacts with the overall naming network both immediately following picture presentation and during articulation (Fig. 8). This is in contrast to Flinker et al. (2015) where trial-averaged interaction measures including AMVAR (Ding et al., 2000) revealed Broca’s area activity only prior to articulation onset, leading to the conclusion that this region exclusively supports articulatory planning and not articulatory execution. While AMVAR also finds no significant interactions for pars triangularis and opercularis during articulation, ARHMM reveals incoming information flow during articulation. This fits best with a feedback mechanism in which each network state terminates activity in the dominant nodes of the prior network state.

**Figure 8:**
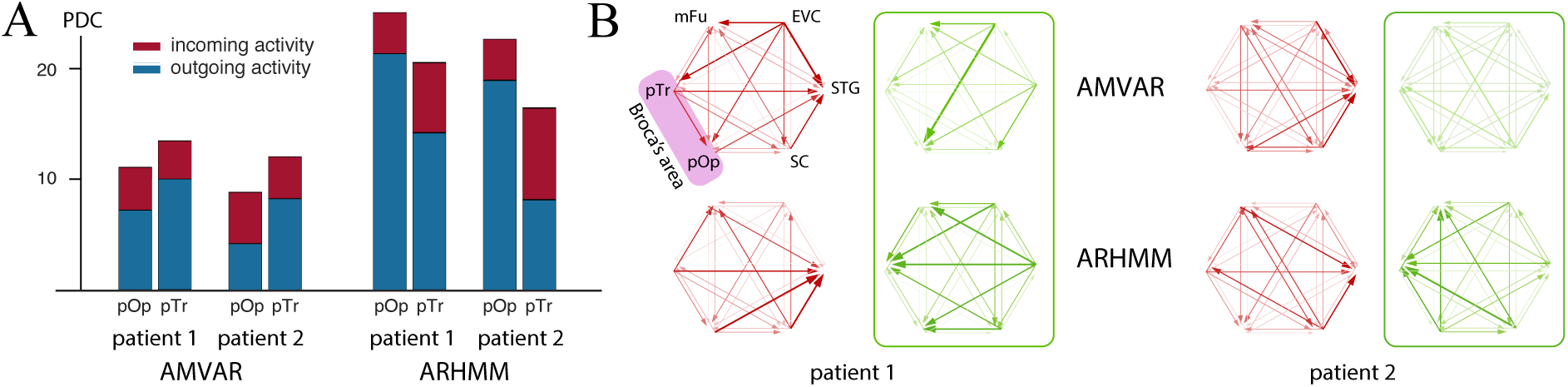
Comparison of AMVAR and ARHMM estimates of activity in Broca’s area during the articulatory state. The ARHMM shows stronger total interactions for than AMVAR analysis, especially for Broca’s area. **A**: Total incoming and outgoing activity (red and blue, respectively) for Broca’s pars opercularis (pOp) and pars triangularis (pTr) during articulation, according to AMVAR and ARHMM models. **B**: Connectivity graphs for AMVAR (top) and ARHMM (bottom), shown for two left-hemispheric patients for the ‘language processing’ (red) and ‘articulation’ state (green).

## 4. Discussion

We identified meaningful cortical states at the single-trial level with a simple but powerful nonlinear model of neuronal interactions — the Autoregressive Hidden Markov Model (ARHMM), a dynamical Bayesian network. We used this model to interpret intracranial recordings at electrodes distributed across the language-dominant hemisphere during a classic language production paradigm, picture naming. The model revealed a consistent progression through three network states that were distinguished by their interaction patterns: visual processing, language processing, and articulation. This analysis of high-resolution intracranial data shows the ARHMM to be a useful tool for parsing network dynamics of human language, and more broadly for quantifying network dynamics during human cognitive function.

### 4.1. Methodological benefits

We have demonstrated that Bayesian dynamical networks can extract structure of dynamical and distributed network interactions from ECoG data. The ARHMM algorithm provides a principled statistical algorithm that can learn brain states in an efficient and unsupervised fashion. Some of the severe limitations of conventional windowed MVAR methods are alleviated by this kind of analysis. One particular virtue is that the ARHMM is sensitive to trial-by-trial timing variations; conventional methods that fail to account for this variability will dilute their estimates of network structure over time, lumping distinct states together or wrongly splitting them. The ARHMM is sensitive to the distributions of network activity, and this allows more refined inferences than the usual *k*-means clustering, which assumes all states have equal, isotropic variability. Bayesian inference of the ARHMM can also incorporate uncertainty, which can be used to determine whether any differences in brain states between subjects, groups, or conditions are significant. While some of these benefits have been previously applied for classifying epileptic activity (Baldassano et al., 2016), or for studying working memory in slow, high-dimensional fMRI signals (Taghia et al., 2018), here we have demonstrated that these models are useful also for discriminating between cognitive states in normal brain function measured at the fast timescales accessible by ECoG measurements. All of these advantages could benefit future work in mining and understanding ECoG data about intact brain computation.

### 4.2. Sequence of states

This work reveals a clear and fairly consistent progression of neural dynamics through three active states in each picture naming trial, which we associate with visual processing, language processing, and finally articulation. These states appear to be meaningful because they are well-aligned with observable behavioral events: picture presentation or articulation. Visual processing was the dominant mode of neural activity for ~250 ms after picture presentation; language processing followed until articulation onset; and articulatory execution lasted for the duration of overt speech production.

The serial progression through these three states is in striking contrast to fully interactive language models, which expect a relatively homogeneous temporal blend of all three states from picture presentation through the end of articulation. Our findings therefore suggest that these states correspond to discrete cognitive processes without strong temporal overlap or interaction (Indefrey and Levelt, 2004).

These cognitive processes — visual processing, language processing, and articulatory execution — are quite broadly defined. In particular, the language processes for speech production are thought to invoke separable cognitive processes supporting semantic, lexical, and phonological elements (Levelt, 1989). Similarly, articulation could be expected to invoke additional stages relating to phono-articulatory loops. The ARHMM did not identify additional distinct states within the language processing interval that might correspond to such elemental processes. If these elemental processes are highly interactive (Dell, 1986), then these elemental processes may effectively blend together into the single state in an ARHMM with recordings at a handful of distributed electrodes. Otherwise, more trials may make it easier to find evidence of such brief processes. The disambiguation of these additional intermediary states is a focus of ongoing work.

### 4.3. Network Interactions

ARHMMs are state-switching models driven by linear, pairwise, directional interactions between network nodes. Consequently, each state is defined by a unique interaction structure. We found that networks in left and right hemispheres had similar structures during visual processing and articulatory execution. Visual processing featured strong interactions between early visual cortex and the rest of the language network. In articulatory execution, interactions were strongest between perisylvian regions: pars triangularis, pars opercularis, subcentral gyrus, and superior temporal gyrus. Pre-articulatory language processing showed a distributed set of interactions across ventral temporal and lateral frontal cortex limited to the language-dominant hemisphere. This evidence is consistent with a bilateral visual processing system (Salmelin et al., 1994) that converges for picture naming to a lateralized language network (Frost et al., 1999), which in turn drives a bilateral articulatory system (Hickok and Poeppel, 2007).

The contrast between interactions during coherent and scrambled naming trials revealed specific cognitive processes supported by discrete sub-networks. Coherent images induced stronger interactions from ventral temporal to lateral frontal regions during visual and language processing, as well as from superior temporal gyrus to the rest of the network during articulation. Scrambled images do not evoke a specific object representation in the brain, requiring only a stereotyped response (“scrambled”). These two functional distinctions between the coherent and scrambled conditions imply that language planning processes are subserved by temporal-to-frontal connections, while phonological motor processes are subserved by frontal-to-temporal connections (Whaley et al., 2016).

### 4.4. Switching linear dynamical systems versus nonlinear dynamics

General nonlinear dynamical systems are too unconstrained to be a viable model for brain activity. Two approaches to constraining the nonlinear dynamics are to use a model that represents smooth nonlinear dependencies, or to use a switching model which combines simpler local representations. Our model is an example of the latter type. One could use hard switches between distinct models, as we and others do (Sahani, 1999; Ghahramani and Hinton, 2000; Fox et al., 2009; Linderman et al., 2017); or one could use smooth interpolations between them (Wang et al., 2006; Yu et al., 2007). Each system will have computational advantages and disadvantages, and it would be beneficial to compare these methods in future work.

### 4.5. Activity-dependent switching versus activity-independent switching, and recognition models versus generative models

Our method is based on a generative model, an assumed Bayesian network that is credited with generating our observed data. One feature of this generative model from Equations 2–3 is latent brain states that transition independently of neural activity. This model therefore cannot generate context- or activity-dependent interactions. Furthermore, the assumed Markov structure enforces exponentially-distributed transitions between network states, which may not reflect the real dynamics of computations. Finally, some of the temporal dynamics of brain states should reflect the interactions’ explicit dependence on sensory input, which is neglected in our model.

There are two properties of our model that mitigate these concerns. First, if multiple latent states produce the same observations, then this could produce non-exponentially-distributed transitions between distinct observable states (Limnios and Oprisan, 2012). Second, even if the model itself corresponds to a *prior* distribution that doesn’t quite have the desired properties, when fit to data the *posterior* distribution can nonetheless exhibit context-dependence and non-exponential transitions between latent states. Essentially, the model’s prior provides enough structure to eliminate many bad fits, while remaining flexible enough to accommodate relevant dynamic neural interactions.

There are many opportunities to generalize the ARHMM to accommodate more desired features. While one always must balance model complexity against data availability, which is typically highly limited for human patients, one can fruitfully gain statistical power by combining data from different subjects to create models with some common behaviors and some individual differences. When applied to common tasks, such models could automatically identify universal properties of neural processing across subjects and even across different tasks.

## Acknowledgements

We thank all the patients who participated in this study; laboratory members at the Tandon lab (Patrick Rollo and Jessica Johnson); neurologists at the Texas Comprehensive Epilepsy Program (Jeremy Slater, Giridhar Kalamangalam, Omotola Hope, Melissa Thomas) who participated in the care of these patients; and all the nurses and technicians in the Epilepsy Monitoring Unit at Memorial Hermann Hospital who helped make this research possible. This work was supported by the National Institute on Deafness and Other Communication Disorders 5R01DC014589-04, the National Institute of Neurological Disorders and Stroke 5U01NS098981-03 and the McNair Foundation.

## Author Contributions

Conceptualization: NT, XP; Methodology: AGS, XP, KJF, NT; Software: AGS; Formal Analysis: AGS; Investigation: AGS, KJF; Data Curation: KJF, NT; Writing — Original Draft: AGS, XP; Writing — Review and Editing: XP, AGS, KJF, NT; Visualization: AGS, KJF, XP; Supervision: XP, NT; Project Administration: XP, NT; Funding Acquisition: NT, XP.

## References

Aertsen A, Bonhoeffer T, Krüger J (1987) Coherent activity in neuronal populations: analysis and interpretation. Physics of cognitive processes pp. 1–34.

Baccalá LA, Sameshima K (2001) Partial directed coherence: a new concept in neural structure determination. Biological Cybernetics 84:463–474.

Baldassano S, Wulsin D, Ung H, Blevins T, Brown MG, Fox E, Litt B (2016) A novel seizure detection algorithm informed by hidden markov model event states. Journal of Neural Engineering 13:036011.

Baum LE, Petrie T, Soules G, Weiss N (1970) A maximization technique occurring in the statistical analysis of probabilistic functions of markov chains. The Annals of Mathematical Statistics 41:164–171.

Conner CR, Chen G, Pieters TA, Tandon N (2014) Category specific spatial dissociations of parallel processes underlying visual naming. Cerebral cortex (New York, N.Y.: 1991) 24:2741–2750.

Conner CR, Ellmore TM, Pieters Ta, Disano Ma, Tandon N (2011) Variability of the relationship between electrophysiology and BOLD-fMRI across cortical regions in humans. The Journal of neuroscience 31:12855–65.

Cox RW (1996) AFNI: Software for analysis and visualization of functional magnetic resonance neuroimages. Comput Biomed Res 29:162–173.

Dale AM, Fischl B, Sereno MI (1999) Cortical surface-based analysis. I. Segmentation and surface reconstruction. NeuroImage 9:179–94.

Dell GS, Schwartz MF, Martin N, Saffran EM, Gagnon Da (1997) Lexical access in aphasic and nonaphasic speakers. Psychological Review 104:801–838.

Dell GS (1986) A spreading-activation theory of retreival in sentence production. Psychological Review 93:283–321.

Dell GS, Burger LK, Svec WR (1997) Language production and serial order: A functional analysis and a model. Psychological Review 104:123.

Dempster AP, Laird NM, Rubin DB (1977) Maximum likelihood from incomplete data via the em algorithm. Journal of the Royal Statistical Society. Series B (Methodological) pp. 1–38.

Ding M, Bressler S, Yang W, Liang H (2000) Short-window spectral analysis of cortical event-related potentials by adaptive multivariate autoregressive modeling: Data preprocessing, model validation, and variability assessment. Biol. Cybern. 83:35–45.

Edwards E, Nagarajan SS, Dalal SS, Canolty RT, Kirsch HE, Barbaro NM, Knight RT (2010) Spatiotemporal imaging of cortical activation during verb generation and picture naming. NeuroImage 50:291–301.

Ellmore TM, Beauchamp MS, Breier JI, Slater JD, Kalamangalam GP, O’Neill TJ, Disano MA, Tandon N (2010) Temporal lobe white matter asymmetry and language laterality in epilepsy patients. NeuroImage 49:2033–2044.

Fisher RA (2006) Statistical methods for research workers, 6th edition. Genesis Publishing Pvt Ltd.

Flinker A, Korzeniewska A, Shestyuk AY, Franaszczuk PJ, Dronkers NF, Knight RT, Crone NE (2015) Re-defining the role of broca’s area in speech. Proceedings of the National Academy of Sciences 112:2871–2875.

Forseth KJ, Kadipasaoglu CM, Conner CR, Hickok G, Knight RT, Tandon N (2018) A lexical semantic hub for heteromodal naming in middle fusiform gyrus. Brain 141:2112–2126.

Fox E, Sudderth EB, Jordan MI, Willsky AS (2009) Nonparametric bayesian learning of switching linear dynamical systems In Advances in Neural Information Processing Systems, pp. 457–464.

Friston KJ, Frith CD, Liddle PF, Frackowiak RSJ (1993) Functional connectivity: The principal-component analysis of large (pet) data sets. Journal of Cerebral Blood Flow & Metabolism 13:5–14.

Fromkin Va (1973) The non-anomalous nature of anomalous utterances. Speech Errors As Linguistic Evidence 47:27–52.

Frost JA, Binder JR, Springer JA, Hammeke TA, Bellgowan PS, Rao SM, Cox RW (1999) Language processing is strongly left lateralized in both sexes: Evidence from functional mri. Brain 122:199–208.

Garrett M (1980) Levels of processing in sentence production Academic Press.

Ghahramani Z, Hinton GE (2000) Variational learning for switching state-space models. Neural computation 12:831–864.

Granger C (1969) Investigating causal relations by econometric models and cross-spectral methods. Econometrica 37:424–38.

Hickok G, Poeppel D (2007) The cortical organization of speech processing. Nature Reviews Neuroscience 8:393.

Horwitz B (1994) Network analysis of cortical visual pathways mapped with pet. Journal of Neurosciences 14:65566.

Indefrey P, Levelt WJM (2004) The spatial and temporal signatures of word production components. Cognition 92:101–144.

Josic K, Rubin J, Matias M, Romo R, editors (2009) Characterizing Oscillatory Cortical Networks with Granger Causality, pp. 169–189 Springer New York, New York, NY.

Kadipasaoglu CM, Baboyan VG, Conner CR, Chen G, Saad ZS, Tandon N (2014) Surface-based mixed effects multilevel analysis of grouped human electrocorticography. NeuroImage 101:215–224.

Kadipasaoglu CM, Conner CR, Whaley ML, Baboyan VG, Tandon N (2016) Category-selectivity in human visual cortex follows cortical topology: A grouped icEEG study. PLoS ONE 11:e0157109.

Kaplan E, Goodglass H WS (1983) The Boston Naming Test Lea and Febiger, Philadelphia.

Korzeniewska A, Crainiceanu CM, Ku R, Franaszczuk PJ, Crone NE (2008) Dynamics of event-related causality in brain electrical activity. Human brain mapping 29 10:1170–92.

Levelt W (1989) Speaking: From Intention to Articulation — Chapter 6 MIT Press.

Limnios N, Oprisan G (2012) Semi-Markov processes and reliability Springer Science & Business Media.

Linderman S, Johnson M, Miller A, Adams R, Blei D, Paninski L (2017) Bayesian Learning and Inference in Recurrent Switching Linear Dynamical Systems In Singh A, Zhu J, editors, Proceedings of the 20th International Conference on Artificial Intelligence and Statistics, Vol. 54 of Proceedings of Machine Learning Research, pp. 914–922, Fort Lauderdale, FL, USA. PMLR.

Morf M, Vieira A, Kailath T (1978) Covariance characterization by partial autocorrelation matrices. Ann. Statist. 6:643–648.

Penny WD, Stephan KE, Mechelli A, Friston KJ (2004) Modelling functional integration: a comparison of structural equation and dynamic causal models. NeuroImage 23:S264–S274.

Pieters TA, Conner CR, Tandon N (2013) Recursive grid partitioning on a cortical surface model: an optimized technique for the localization of implanted subdural electrodes. Journal of Neurosurgery 118:1086–1097.

Rajapakse JC, Zhou J (2007) Learning effective brain connectivity with dynamic bayesian networks. NeuroImage 37:749–760.

Rapp B, Goldrick M (2000) Discreteness and interactivity in spoken word production. Psychological Review 107:460–499.

Sahani M (1999) Latent variable models for neural data analysis California Institute of Technology Pasadena, CA.

Sahin NT, Pinker S, Cash SS, Schomer D, Halgren E (2009) Sequential processing of lexical, grammatical, and phonological information within Broca’s area. Science 326:445–9.

Salmelin R, Hari R, Lounasmaa O, Sams M (1994) Dynamics of brain activation during picture naming. Nature 368:463.

Schwarz G (1978) Estimating the dimension of a model. Ann. Statist. 6:461–464.

Snodgrass JG, Vanderwart M (1980) A standardized set of 260 pictures: Norms for name agreement, image agreement, familiarity, and visual complexity. Journal of Experimental Psychology: Human Learning & Memory 6:174–215.

Taghia J, Cai W, Ryali S, Kochalka J, Nicholas J, Chen T, Menon V (2018) Uncovering hidden brain state dynamics that regulate performance and decision-making during cognition. Nature communications 9:2505.

Tandon N (2008) Cortical mapping by electrical stimulation of subdural electrodes: language areas In Luders H, editor, Textbook of Epilepsy Surgery, chapter 109, pp. 1001–1015. McGraw Hill.

Wada J, Rasmussen T (2007) Intracarotid Injection of Sodium Amytal for the Lateralization of Cerebral Speech Dominance. Journal of Neurosurgery 106:1117–1133.

Wang J, Hertzmann A, Fleet DJ (2006) Gaussian process dynamical models In Advances in Neural Information Processing Systems, pp. 1441–1448.

Whaley ML, Kadipasaoglu CM, Cox SJ, Tandon N (2016) Modulation of orthographic decoding by frontal cortex. Journal of Neuroscience 36:1173–1184.

Wulsin DF, Fox EB, Litt B (2014) Modeling the complex dynamics and changing correlations of epileptic events. Artificial Intelligence 216:55–75.

Yu BM, Kemere C, Santhanam G, Afshar A, Ryu SI, Meng TH, Sahani M, Shenoy KV (2007) Mixture of trajectory models for neural decoding of goal-directed movements. Journal of Neurophysiology 97:3763–3780.

Zheng X, Rajapakse JC (2006) Learning functional structure from fmr images. NeuroImage 31:1601–1613.

